# phylobar: an R package for multiresolution compositional barplots in omics studies

**DOI:** 10.1101/2025.11.05.686662

**Authors:** Megan Kuo, Kim-Anh Lê Cao, Saritha Kodikara, Jiadong Mao, Kris Sankaran

## Abstract

**Summary:** Stacked barplots, though widely used in microbiome studies, can obscure important patterns in microbiome data. They omit rare taxa and can mask shifts that emerge at finer taxonomic levels. To address this issue, we introduce phylobar, an R package that interactively links stacked barplots with overview phylogenetic or taxonomic hierarchies. The interface allows users to collapse or expand subtrees, paint color palettes interactively, and search for specific taxa. This allows comparison across taxonomic resolutions that are hidden in static overviews. phylobar works with any omics data with hierarchical organization, including cell type hierarchies, as we demonstrate in a case study of immune cell composition in COVID-19 patients.

**Availability and Implementation:** phylobar is available as an R package on GitHub. It uses the htmlwidgets library to link interactive D3 visualizations with R. The interactive plots can be embedded within R Markdown or Quarto notebooks, and views can be exported as vector graphics files. The package is open source and documented at https://mkdiro-O.github.io/phylobar.

## Introduction

Stacked barplots are widely used to visualize microbiome community structure. These plots divide each bar – representing a single sample – into colored segments. The colors denote taxonomic categories; the heights reflect group frequencies. This provides a useful exploratory overview for understanding the main taxonomic groups that are present across the data set. The view also supports comparisons in community structure across samples, especially when the samples are ordered in a meaningful way—for example, sorting by time or sample similarities. These plots have driven biological discovery in the microbiome literature. For example, they have helped group samples into community state types [7], highlight heterogeneity among dysbiotic samples [24], and clarify response to environmental interventions [19]. Several R packages are now available to generate these plots [16, 1, 29, 11]. They can be made from community composition matrices or directly from microbiome-specific data structures, like those exported by phyloseq [18] or TreeSummarizedExperiment [10] objects.

Stacked barplots use color to graphically encode taxonomic categories. This imposes a basic constraint – when the color palette grows, distinctions become more and more challenging to make. Researchers typically solve this in one of two ways: visualize only high-level taxonomic categories, or show low-level categories for abundant groups and aggregate rare taxa into a bin for “Other” [28]. Yet some widely used packages demonstrate this problem in their own documentation. The qiime2 viewer uses 200 colors in a single bar plot [4], and an example in MicrobiomeStat uses 43 [2]. These palettes make the plots extremely difficult to use.

Recently, the StackbarExtend package [6] addressed this problem by assigning similar colors to related taxonomic categories. In this way, color indicates taxonomic assignment while ensuring broader categories remain distinguishable. We also use taxonomy to guide the stacked bar plot. However, instead of color gradients, phylobar uses interactivity. Specifically, phylobar exploits hierarchical structure and the focus-plus-context principle [21, 8, 20], allowing users to collapse and expand the hierarchy to different resolutions and paint the barplot by hovering the mouse over different subtrees. Hence, users can interactively test different color palettes and identify taxonomic groups with interesting variation before finalizing the visualization. Further, this approach allows direct comparison of multiple levels of taxonomic resolution, since large and small subtrees can be painted simultaneously.

phylobar applies to compositional visualization in other hierarchically structured omics data. This includes pseudobulk analysis of single-cell data, since cell types can be organized into nested categories across resolution levels. Example 2 below demonstrates phylobar’s utility in this single-cell setting.

In addition to initial exploratory analysis, phylobar also enables post-modeling exploration. For example, when a model identifies differentially abundant taxa, the same interactive barplot can be used to visualize the distribution and variability of those taxa across samples and groups, providing a clear, sample-level view of model-derived findings. This approach preserves the benefits of the original stacked bar plot, allowing overviews of the entire data set and comparison across taxonomic categories. But viewers can now interactively browse taxonomic groups (including rare ones), no longer needing to accept a static set of coarse or pre-defined categories.

### Overview of package functionality

The main visualization function in the phylobar package is called `phylobar`. This function has two required inputs, (1) a data matrix of abundances with samples in rows and taxa in columns, and (2) a phylo structure defining the taxonomic hierarchy. The leaf nodes of the phylo must match the column (taxa) names of the sample abundance matrix. A leaf node is the lowest level of the tree hierarchy and represents the finest level of taxonomic resolution. Sample abundances need not sum to one, and users can provide non-compositional normalizations to phylobar, resulting in stacked bar plots where the heights across bars are not necessarily equal. phylobar requires a phylo object representing the tree. Considering that hierarchical information often exists in less structured formats, like taxonomic tables, the package includes a function (`taxonomy_to_tree`) that converts these formats into the required phylo structure. Note that the resulting tree does not have to be a binary tree; that is, each internal node is not restricted to having only two child nodes.

phylobar includes two panels (Fig 1a). The left panel shows a taxonomic hierarchy drawn as a tree. The right panel shows a stacked barplot where each rectangle within a sample’s bar is corresponds to one leaf from the tree. Samples can be ordered either by hierarchical clustering or by their input data matrix rows. The latter option enables users to reorder samples based on predefined characteristics, such as chronological time, experimental group, or other relevant study factors.

**Fig. 1.**
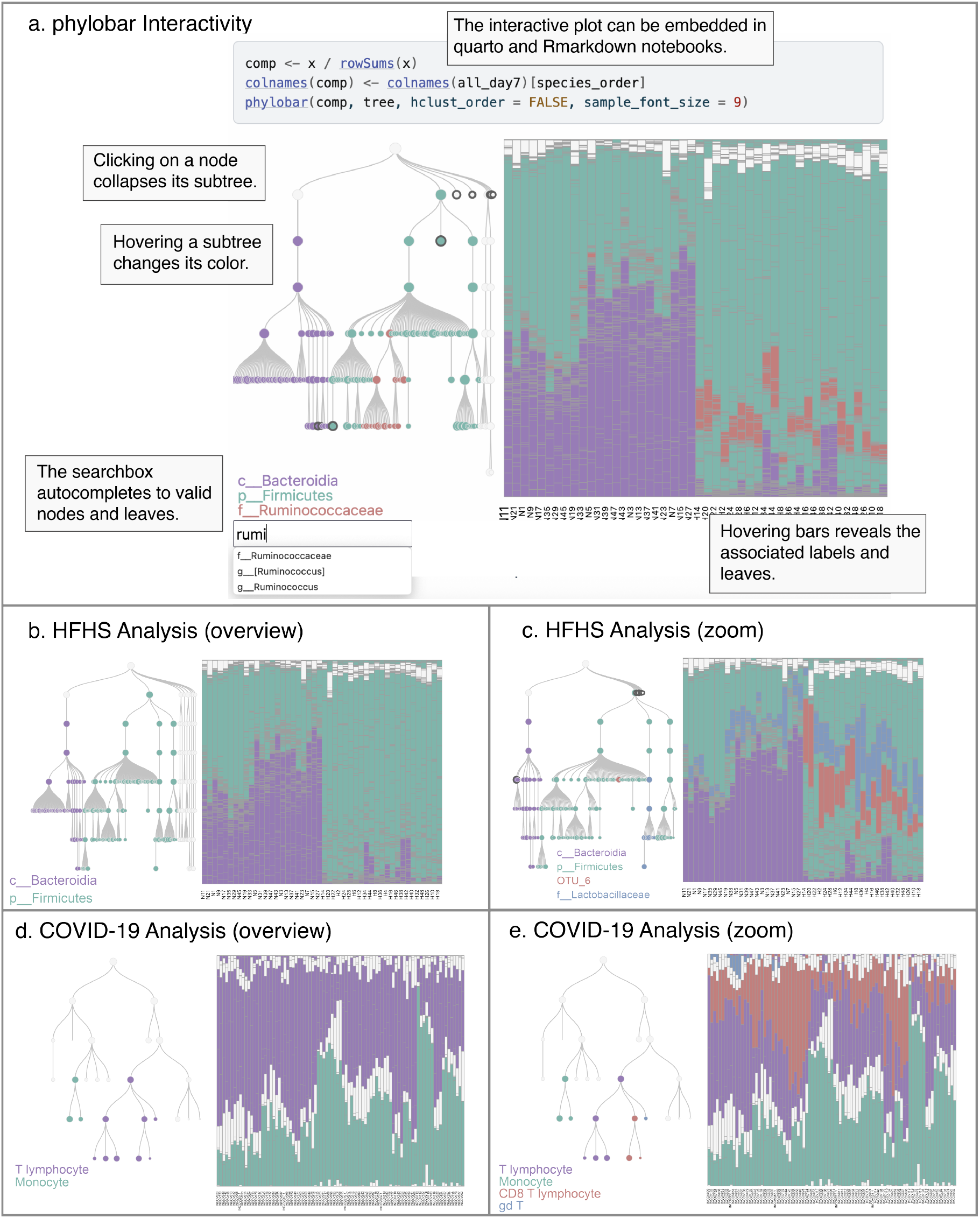
(a) An overview of phylobar’s display and interactivity. Clicking or hovering over tree nodes and barplot compositions helps focus comparisons across taxa. A search box supports lookup of individual taxa. (b) High-level overviews confirm Bacteroidia and Firmicutes shifts under the HFHS intervention from Susin et al. (2020) [23]. (c) Collapsing unrelated subtrees focuses the view on these two groups. Searching for family Lactobacillaceae confirms its HFHS association, and hovering over bars reveals the elevated abundance of OTU 6 in the HFHS group. d. An immune cell type hierarchy coupled with cell type compositions from a study of immune factors in COVID-19 severity. e. Hovering over nodes for T cell subtypes replaces the purple stacked bar portions from panel (d) with colors associated with the CD8 T lymphocyte and *γδ* subtypes.

Hovering near a node paints the associated subtree with the user-provided color palette and highlights the associated leaves in the stacked bar plot. Pressing the Control key fixes the currently painted subtrees and adds a new color for painting. Colors can be added until the initial palette is exhausted. By default, the initial palette is 6 colors. Once the palette is exhausted, Control cycles through previously used colors. Alternatively, the number keys select specific colors in the palette. While painting, a color legend is interactively updated to display each highlighted subtree’s root label. This tree painting strategy makes it possible to compare the abundance of taxonomic categories, some of which may be rare or at different levels of taxonomic resolution, across the full collection of samples in the data.

Large trees benefit from trimming taxonomic subgroups of limited interest. To this end, users can click on a to collapse all of its descendants. The collapsed node appears with a thicker border to indicate the hidden subtree. The entire tree layout is redrawn to make use of the newly freed space. When a subtree is collapsed, the associated leaf nodes in the stacked bar plot are merged into a single rectangle corresponding to the entire subtree. If a color had been present only within the descendants of a collapsed subtree, that color is removed from the palette. Clicking the the collapsed subtree expands it to its previous state.

Hovering over a bar in the stacked barplot magnifies its sample name. When bars are tightly packed, labels can optionally be hidden until the sample is hovered over. Similarly, when a dataset includes many samples, plotting all samples becomes impractical. In this case, the package can select representatives from hierarchical clusters derived from the data set, and the stacked bar plot will show one representative from each cluster.

Altogether, this interactivity supports efficient comparison of taxonomic categories across subtrees. The visualization’s appearance is customizable, and users can modify color palettes, font sizes, and component dimensions. Labels can be placed directly on subtrees rather than in a separate legend. If the analyst finds a useful view, then pressing the Escape key freezes the view for export into a Scalable Vector Graphics (SVG) file. Pressing escape again resumes the interaction. While simple, phylobar supports exploratory analysis of microbiome community structure at different levels of resolution, preserving distinctions even at strain-level variation.

## Methods/Implementation

phylobar is an R package whose visualizations are built using the D3 JavaScript library [5]. D3 transforms JavaScript data arrays into graphical layout coordinates and provides methods for Document Object Model (DOM) manipulation. phylobar uses d3.hierarchy and d3.stack for tree and stacked bar plot visualizations, respectively. The resulting hierarchical layout includes methods for enumerating children, descendants, and leaf nodes, which support subtree painting and collapsing. A Delaunay triangulation around the tree nodes triggers mouseover events when the cursor moves between node neighborhoods, allowing smoother interaction than direct node manipulation. In response to mousemoves and clicks, D3 manipulates the DOM to change subtree and barplot colors and resolution. The DOM is modified in place without re-rendering the entire view. For example, hovering a subtree updates fill attributes in DOM elements without restructuring them. This ensures smooth response to user inputs and is a key advantage of browser-based visualization. A standalone JavaScript package, phylobar-js, wraps the core data manipulation and visualization functions, and is available on NPM [3]. This package exports reusable components for building interactive hierarchical visualizations, including functions for interactively collapsing and expanding trees, searching for string matches across hierarchy labels, and detecting mouse hover events across node neighborhoods.

The htmlwidgets R package [26] bridges phylobar’s JavaScript implementation and R interface. This package converts R matrices and lists into JavaScript arrays and objects. htmlwidgets also creates an SVG canvas for D3 visualizations. This canvas can be embedded within Quarto [17] or Rmarkdown [9] notebooks with multiple visualizations per notebook. Views can also be exported to scalable vector graphics files, allowing customization (e.g., font families and edge stroke types) in vector graphics software like Illustrator or Inkscape.

## Results

### Example 1: High-fat high-sugar diet and the gut microbiome

The gut microbiome is known to be reshaped by high-fat, high-sugar (HFHS) diets. Susin et al. (2020) [23] were the first to analyze the HFHS mouse model to characterize diet-driven microbiome effects using samples collected by Lê Cao at the University of Queensland Diamantina Institute. They randomized 47 C57BL/6 female mice into HFHS and control diet groups and gathered fecal samples at days 0, 1, 4, and 7. We use phylobar to re-analyze the preprocessed 16S rRNA sequencing data from Kodikara et al. (2025) [13]. These data were generated using Illumina MiSeq and taxnomic representation was determined using QIIME 1.9.0. To concentrate on long-term effects of the diet intervention, we filter to samples from day 7. In our stacked barplot, we place all control and HFHS samples on the left and right, respectively. Within the two treatment groups, we apply complete link hierarchical clustering on the euclidean distance between community composition vectors and sort samples according to the leaf order on the resulting trees. We first check whether we observe an effect on the Bacteroides vs. Firmicutes ratio, which has previously been found to be associated with HFHS diet. Fig 1b shows the stacked barplots when hovering over class Bacteroida and Phylum Firmicutes. Consistent with earlier studies, Bacteroidia appears more abundant in the control samples, while Firmicutes is elevated in the HFHS condition. Next we asked whether we could decompose this observation into finer effects within narrower taxonomic groups. Fig 1c shows the visualization after highlighting a particular strain, OTU 6, and searching for family Lactobacillaceae, both of which are members of phylum Firmicutes. Subtrees unrelated to Bacteroidia and Firmicutes have been collapsed. Consistent with the previous view, abundances of both groups increase in the HFHS condition, but they change in qualitatively different ways. OTU 6 is absent in all but a handful of control samples, but it is one of the most abundant taxa in the HFHS communities. In contrast, family Lactobacillaceae is present in most control samples, but at lower levels than in the HFHS condition. In this example, all three forms of interaction–browsing, sorting, and searching– proved important. Interactivity makes it straightforward to compare samples across multiple levels of taxonomic resolution and helps avoid the aggregation required in static alternatives. This encourages multiresolution exploration and helps generate precise, biologically relevant hypotheses.

### Example 2: Immune cell type composition in mild to severe COVID-19 patients

In addition to microbiome data, we can use phylobar to visualize other types of compositions. We applied phylobar to a single-cell RNA-seq dataset from Su et al. (2020) [22] to examine how immune cell type composition changes across mild, moderate, and severe COVID-19. The input was the proportions of immune cell types from 120 COVID-19 patients’ blood samples. Patients were first grouped by their disease severity and then ordered by hierarchical clustering, so that patients with similar cell type compositions within each severity group were more closely placed. A cell type tree representing the hierarchical structure of immune cells was manually defined.

Based on the phylobar visualization we observed that T cells and monocytes made up the majority of cellular populations in patient samples. We also observed an overall increase of monocytes and decrease of NK and T cells (Fig 1d), especially CD8 T cells, in moderate and severe COVID-19 patients. These changes were also highlighted in Su et al. (2020) [22]. Another feature revealed by phylobar that was not characterized in Su et al. (2020) [22] was the presence of *γδ* (gd in the cell type tree) T cells in milder COVID-19 patients exclusively (Fig 1e). This finding echos some more recent works suggesting that *γδ* T cells may play an important role in the immune response to the pathogenic SARS-Cov-2 virus [25, 27].

## Discussion

phylobar adapts the focus-plus-context principle to compare microbiome sample composition across taxonomic groups and levels of resolution. The package interactively links hierarchical overviews with barplots of sample composition in selected subtrees. This approach solves a longstanding limitation of stacked barplots when applied to omics data visualization, where the number of features exceeds what can be meaningfully represented with a qualitative color palette. The display requires only two inputs, allowing for its broad applicability. The package documentation includes nine vignettes: five data analysis walkthroughs (including those that reproduce the microbiome and single-cell examples presented above), and four guides on preparing package inputs and refining visual outputs. For future work, we note that specialized interactions could be tailored to specific experimental designs, like the longitudinal data in [13] or pseudotime data in [14]. We also note that phylobar can be adapted to hierarchies that are learned from sample abundance or expression profiles [12, 15], rather than specified in advance. Code and documentation are available at https://mkdiro-O.github.io/phylobar.

